# VAPOR: Variational autoencoder with transport operators decouples co-occurring biological processes in development

**DOI:** 10.1101/2024.10.27.620534

**Authors:** Jie Sheng, Daifeng Wang

## Abstract

**Background:** Emerging single-cell and spatial transcriptomic data enable the investigation of gene expression dynamics of various biological processes, especially for development. To this end, existing computational methods typically infer trajectories that sequentially order cells for revealing gene expression changes in development, e.g., to assign a pseudotime to each cell indicating the ordering. However, these trajectories can aggregate different biological processes that cells undergo simultaneously–such as maturation for specialized function and differentiation into specific cell types–which do not occur on the same timescale. Therefore, a single pseudotime axis may not distinguish gene expression dynamics from co-occurring processes.

**Methods:** We introduce a method, VAPOR (variational autoencoder with transport operators), to decouple dynamic patterns from developmental gene expression data. Particularly, VAPOR learns a latent space for gene expression dynamics and decomposes the space into multiple subspaces. The dynamics on each subspace are governed by an ordinary differential equation model, attempting to recapitulate specific biological processes. Furthermore, we can infer the process-specific pseudotimes, revealing multifaceted timescales of distinct processes in which cells may simultaneously be involved during development.

**Results:** Initially tested on simulated datasets, VAPOR effectively recovered the topology and decoupled distinct dynamic patterns in the data. We then applied VAPOR to a developmental human brain scRNA-seq dataset across postconceptional weeks and identified gene expression dynamics for several key processes, such as differentiation and maturation. Moreover, our benchmarking analyses also demonstrated the outperformance of VAPOR over other methods. Additionally, we applied VAPOR to spatial transcriptomics data in the human dorsolateral prefrontal cortex. VAPOR captured the ‘inside-out’ pattern across cortical layers, potentially revealing how layers were formed, characterized by their gene expression dynamics.

**Conclusion:** VAPOR is open source for general use (https://github.com/daifengwanglab/VAPOR) to parameterize and infer developmental gene expression dynamics. It can be further extended for other single-cell and spatial omics such as chromatin accessibility to reveal developmental epigenomic dynamics.

## 1 Introduction

Development is complex, orchestrated by multiple, interrelated biological processes that occur simultaneously[1, 2, 3]. These processes include ubiquitous ones like cellular proliferation, as well as system-specific processes like differentiation, migration, or functional maturation, each occurring at varying timescales and rates[4, 5, 6, 7]. These processes are fundamentally driven by dynamic changes in gene expression at the individual cell level. Recent advancements in single-cell and spatial genomics technologies have provided unprecedented resolution in capturing these gene expression dynamics during development. For instance, single-cell RNA sequencing (scRNA-seq) can capture snapshots of gene expression profiles from individual cells at different stages, showing gene expression changes over time[8, 9, 10, 11]; while spatial transcriptomics (ST) measures gene expression from cells within their tissue context, showing how gene expression varies across different regions within a tissue section[12, 13, 14]. These technologies provide researchers with distinct aspects of gene expression dynamics during development - the temporal and spatial context, respectively. However, the interpretation of this high-dimensional, heterogeneous data remains challenging, particularly in decoupling the gene expression dynamics underlying multiple co-occurring biological processes.

Many computational methods have been developed to analyze single-cell and spatial transcriptomics data in development. However, they often provide aggregated views of these processes, capturing overall trends of development but potentially overlooking the contributions of individual biological processes. For instance, graph-based trajectory inference methods [15, 16, 17] map cells onto graph or tree-like trajectories and assign a pseudotime to represent the temporal ordering of cells during development. However, relying on a single pseudotime axis oversimplifies development by aggregating multiple, co-occurring processes onto a unified scale, which obscures the unique contributions of each process to the complex gene expression dynamics. Another type of approach is RNA velocity methods[18, 19, 20, 21] that focus on higher-resolution gene expression dynamics by modeling splicing kinetics using ordinary differential equations (ODEs). Unlike trajectory inference methods that order cells along developmental paths, RNA velocity methods capture immediate transcriptional changes and predicts short-term gene expression of cells. However, RNA velocity methods typically assume a uniform rate of gene activity change (known as kinetics rate). This assumption may not hold as genes can exhibit multiple distinct kinetic regimes during development due to the various processes involved [22].

Recent advancements have attempted to address some of the limitations. Multifaceted analysis methods, such as TopicVelo[23] and GeneTrajectory[24], aim to decompose complex gene expression dynamics and separate co-occurring processes from gene expression data. However, these methods still have limitations. TopicVelo[23] is an RNA velocity-based method, thus limited in analyzing the biological processes where splicing information is unavailable or less relevant, such as spatial transcriptomics or single-nuclei RNA sequencing. Additionally, the requirement for a predetermined number of processes also constrains its ability to discover novel and unexpected processes. GeneTrajectory[24] focuses on genes by partitioning them into sets associated with independent components of gene expression dynamics. However, at the cellular level, it remains unclear how these components relate t o specific cell populations, limiting our understanding of how co-occurring processes independently or cooperatively shape cell development.

To address those challenges, we develop a method, variational autoencoder[25] with transport operators[26] (VAPOR), to decouple gene expression dynamics and identify distinct biological processes. Initially tested on simulated datasets, VAPOR effectively decoupled distinct dynamic patterns and recovered the topology in the data. We then applied VAPOR to a developmental human brain scRNA-seq dataset[8] during postconceptional weeks and identified gene expression dynamics for two key developmental processes, differentiation and cell-type-specific maturation. Additionally, we applied VAPOR to spatial transcriptomics data in the human dorsolateral prefrontal cortex[12]. Our model captured the preserved ‘inside-out’ pattern across cortical layers, revealing how gene expression dynamic patterns align with layer formation.

## 2 Methods

VAPOR combines a variational autoencoder[25] (VAE) with transport operators[26] (TO) to learn a latent representation of cells that captures gene expression dynamics associated with distinct biological processes (**Fig. 1**). The VAE (**Fig. 1a**) projects high-dimensional gene expression data into a lower-dimensional latent space while persevering essential dynamics. The TO (**Fig. 1b**) models the dynamics of the latent mean parameters *μ*, decomposing the latent space into multiple subspaces, each corresponding to a distinct biological process. The dynamics in each subspace are governed by ordinary differential equations (ODEs), modeling processes occurring simultaneously but potentially at different timescales and rates. Consequently, we infer process-specific pseudotimes (**Fig. 1c**), allowing for a multifaceted ordering of cells across distinct biological processes and providing a detailed view of how cells progress through these processes. Pseudocode of VAPOR can be seen in **Algorithm** 1.

**Figure 1:**
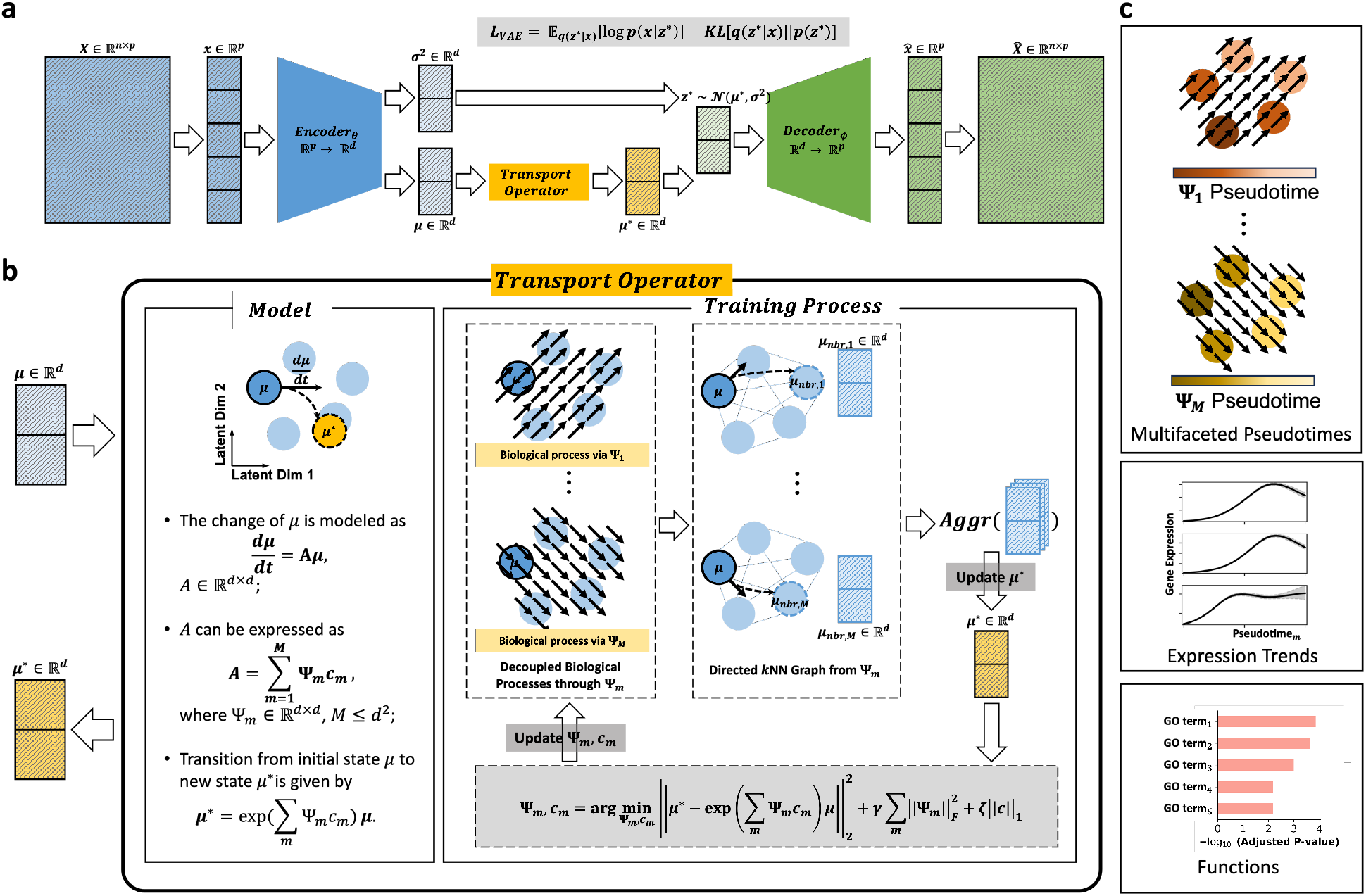
Overview of VAPOR. a, Schematic of VAPOR’s architecture. The input gene expression matrix is encoded into a lower-dimensional probabilistic latent space, represented by a multivariate Gaussian distribution 𝒩 ∼ (*μ, σ*^2^). The transport operator transforms the mean parameter *μ* to *μ*^*^, modeling latent dynamics associated with different biological processes. Then, the transformed mean parameter *μ*^*^ defines the latent distribution used to generate the latent variable *z*^*^. Finally, *z*^*^ is decoded back to reconstruct the gene expression data. **b, Transport Operator model and training process**. Changes in the latent space are modeled by the ODEs applied to *μ*. Without ground truth for the actual new states, we employ an EM-like iterative approach. For each cell, a “new state” is sampled from its *k*-nearest neighbors for each decoupled process Ψ_*m*_. These new states are aggregated to form a single representative “new state”, which is used to optimize Ψ_*m*_ and *c*_*m*_ by minimizing the loss function. The updated Ψ_*m*_ then guides the selection of “new states” through a directed *k*-NN graph in the next iteration. **c, Decoupled** Ψ**s recapitulate distinct biological processes**. The learned model allows for multifaceted analysis of co-occurring biological processes. 1) Process-specific pseudotime: cells are assigned pseudotime for each process, captured by Ψ_*m*_, showing their progression along distinct processes; 2) Gene expression dynamics: the change in gene expression levels across pseudotime for each process allows identification of process-specific gene expression patterns; 3) Functional annotation: gene ontology (GO) analysis on the genes associated with each decoupled process reveals the biological functions and pathways in different aspects of development.

### 2.1 Variational Autoencoder Preserves Gene Expression Dynamics in Latent Space

We utilize a VAE to capture the dynamics of high-dimensional gene expression data. The encoder *E*(·) maps the input gene expression matrix *X* ∈ ℝ^*n×p*^, where *n* is the number of cells and *p* is the number of genes, into a *d*-dimensional latent space. The latent space is parameterized as a multivariate Gaussian distribution 𝒩 (*μ*, diag(*σ*^2^)) with mean *μ* ∈ ℝ^*d*^ and variance *σ*^2^ ∈ ℝ^*d*^.

To model the preserved dynamics, we introduce a transport operator *TO*(·) (details in **Section 2.2**) that transforms the latent mean *μ* to a new state *μ*^*^,

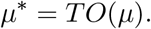

We apply transport operator on *μ* instead of *σ* as considering *μ* represents the central tendency of the latent state, which could be interpreted as the “average” or “most likely” state of a cell; while *σ* represents the uncertainty or variability around that state. By not transforming, we’re assuming that the uncertainty structure remains relatively invariant during transitions.

The decoder *D*(·) then reconstructs the original gene expression data from the latent variables

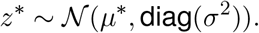

#### Algorithm 1

VAPOR: Variational autoencoder with transport operators.

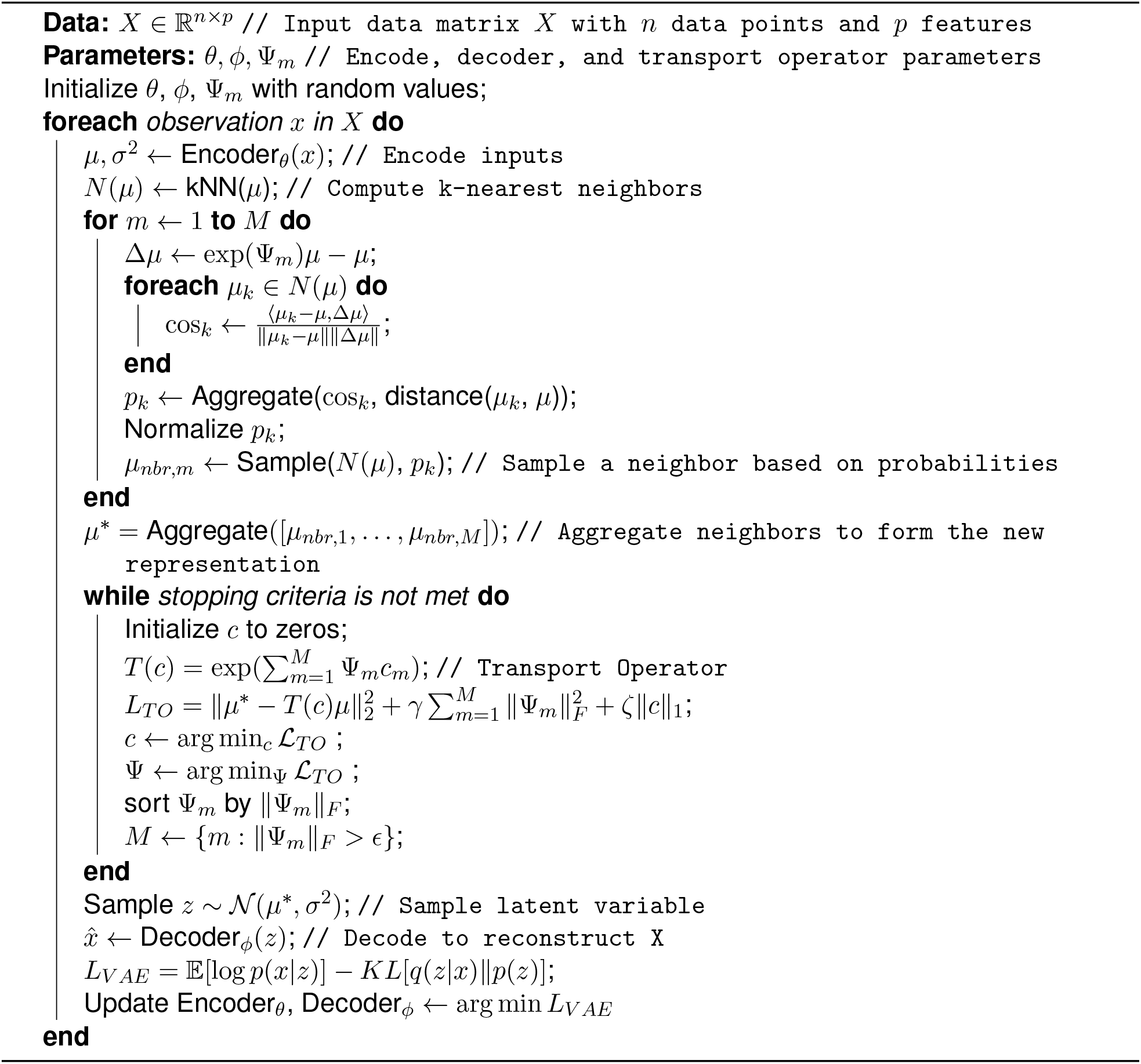

To learn the latent space, VAE minimizes the following loss function,

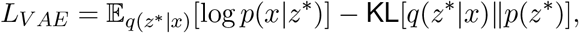

where *q*(*z*^*^|*x*) is the approximate posterior distribution of the latent variable given the data; *p*(*z*^*^) is the prior distribution over the latent variables; and KL[·||·] denotes the Kullback-Leibler divergence.

### 2.2 Transport Operators Decouples Latent Dynamics into Distinct Processes

#### Modeling

VAPOR extends the VAE model by applying a transport operator (TO)[26] to the latent mean *μ*, modeling changes corresponding to distinct biological processes. The change in the latent mean *μ* is modeled using an ordinary differential equation (ODE):

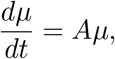

where *A* ∈ ℝ^*d×d*^ is a learnable matrix representing the dynamic driving changes in *μ*. Rather than directly learning *A*, VAPOR decomposes it into a weighted sum of basis matrices Ψ_*m*_ ∈ ℝ^*d×d*^, that is

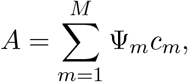

where each Ψ_*m*_ corresponds to a different biological process; *c*_*m*_ is a scalar coefficient representing the contribution of the *m*-th process; and *M* is the predetermined number of processes we aim to decouple, *M* ≪ *d*^2^.

The transport operator moves *μ* to a new state *μ*^*^ (assuming *t* = 1) by

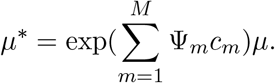

#### Learning TO

Without ground truth for the actual new states, we employed an EM-like iterative approach (Fig 1b) to learn the transport operators. The workflow to the algorithm is as follows:

##### Step 1, Initialization

We initialize Ψ_*m*_ matrices with random values and *c*_*m*_ with zeros, where *m* = 1, · · ·, *M*.

##### Step 2, Expectation step

For each cell *i*, we select “new states” from its *k*-nearest neighbors, one for each Ψ_*m*_. The selection is guided by the current estimates of Ψ_*m*_, favoring neighbors that align well with the direction of change

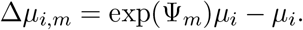

That is, we prefer neighbors that are consistent with the expected progression along each process Ψ_*m*_.

##### Step 3, Aggregation multiple processes

Aggregate the multiple “new states” for each cell *I* (one per process) to form a single representative “new state” 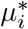:

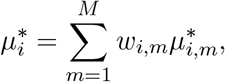

where *w*_*m*_ are weights learned from the VAE updating process. The VAE determines *w*_*i,m*_ based on how well each “new state” 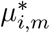 contributes to reconstsructing the original gene expression data, implying the informativeness of each process for that cell.

##### Step 4, Maximization step

Using these aggregated “new states”, we optimize Ψ_*m*_ and *c*_*m*_ by minimizing the loss function:

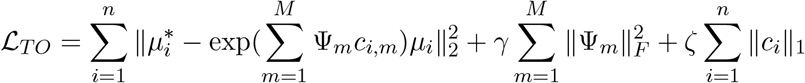

where *γ* and *ζ* are regularization parameters, ∥ · ∥_*F*_ denotes the Frobenius norm, and ∥ · ∥_1_ is the *L*1 norm encouraging sparsity in the coefficients.

##### Step 5, Sorting and Pruning of Processes

After each iteration, we refine our set of processes Ψ_*m*_. First, we sort the processes based on their Frobenius norm ∥Ψ_*m*_∥_*F*_ in descending order. This ordering reflects the relative importance of each process. Then, we apply a filtering step:

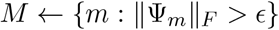

where *ϵ* is a small threshold value, which removes negligible impact processes. This steps help us retain the the most significant processes.

##### Iteration

Above step 2-5 are repeated for a fixed number of iterations or until convergence criteria are met.

### 2.3 Training Process of VAPOR

In VAPOR, we alternate between training the VAE and TO in separate phases. We first optimize the VAE for several epochs to obtain a meaningful latent representation that captures the underlying dynamics of the gene expression. This is achieved by optimizing *L*_*V AE*_. With the VAE parameters fixed, we then train the TO by optimizing *L*_*T O*_. After training the TO, we update the VAE by re-training it using the update *TO*(·), which helps the VAE to incorporate the dynamics captured by the *TO*(·). We repeat this alternating training process until convergence.

### 2.4 Process-Specific Operators Ψ_*m*_ Reveals Distinct Biological Processes

VAPOR identify decoupled co-occurring biological processes through the process-specific operators Ψ_*m*_, and estimate multifaceted pseudotime for each process.

#### 2.4.1 Computing Multifaceted Pseudotimes

To represent the progression of cells along each biological process, we estimate pseudotime based on its corresponding operator Ψ_*m*_. We computes a velocity-like vector analogous to RNA velocity for each cell *i*,

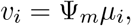

where *μ*_*i*_ is the cell’s latent mean vector. We utilize the `scvelo.tl.velocity_pseudotime()` function from scVelo[19] to compute pseudotimes for each Ψ_*m*_.

#### 2.4.2 Visualization of decoupled processes

To visualize each process, we adapt the velocity graph approach from scVelo[19]. While scVelo computes transition probabilites using cosine similarities between RNA velocity and potential state transitions in terms of gene expression (*x*_*j*_ − *x*_*i*_), we compute similarities between

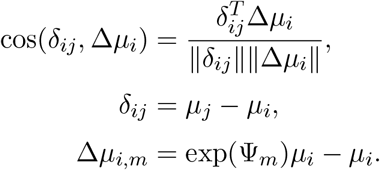

Here, exp(Ψ_*m*_)*μ*_*i*_ represents the predicted future state of cell *i* under process *m* in latent space, and Δ*μ*_*i,m*_ captures the expected change in the cell’s state in latent space. We the use the `scvelo.tl.velocity_graph()` function from scVelo[19] for visualization.

#### 2.3.3 Investigating Gene Expression along Pseudotimes

To understand how gene expression changes along each process-specific pseudotime, we analyze the expression trends of genes across the pseudotimes for each Ψ_*m*_. We use generalized additive models (GAMs) (by Python `pyGLM` library) to model the relationships between gene expression and pseudotime. We plot the fitted expression curves against pseudotime for selected genes.

#### 2.4.4 GO enrichment analysis

To gain biological insights into the processes captured by each Ψ_*m*_, we perform Gene Ontology (GO) enrichment analysis on genes associated with each process. We compute the Pearson correlation coefficients between gene expression and the computed pseudotime for each process *m*. We rank the genes based on the absolute value of their correlation coefficients, selecting the top genes that are most positively and negatively correlated with pseudotime. Using the top-ranked genes for each process, we conduct GO enrichment analysis (utilizing Python `gseapy` library against the `‘GO_Biolgocial_Process_2023’` database).

### 2.5 Datasets and Data Preprocessing

#### 2.5.1 Simulation

We followed GeneTrajectory’s[24] simulation approach. Basically, for linear process, we first assigned each cell a pseudotime *t*_*u*_ uniformly distributed in the range [0, *T*]. Each gene *i* was assigned a peak expression time 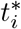, also uniformly distributed in the range [0, *T*]. The expected expression of gene *i* in cell *u* is normally distributed, centered at 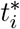, given by

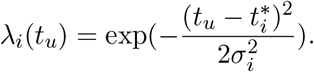

Additionally, to simulate experimental noise, we simulated the observed gene expression counts from a Poisson distribution with mean *λ*_*i*_(*t*_*u*_). For cyclic process, we just need to modify *λ*_*i*_(*t*_*u*_) as

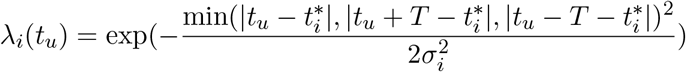

to account for periodic patterns.

We simulated three scenarios: (1) a linear progression where genes sequentially reach their peak expression; (2) a cyclic process where genes show periodic expression patterns over time; and (3) a combination of linear and cyclic processes, where each cell state is determined by two independent variables: a pseudotime along the linear process and a pseudotime along a cyclic process.

#### 2.5.2 scRNA-seq of Human cerebral cortex

The raw data set is from [8] and is available in the Gene Expression Omnibus repository under accession number GSE162170. We included scRNA-seq data from three experimental time points: post-conceptional week (PCW) 20, PCW21, and PCW24. The count matrix are normalized. The top 2,000 highly variable genes are selected. Then the normalized matrix is scaled to unit variance and zero mean for each features (genes).

#### 2.5.3 ST data of human dorsolateral prefrontal cortex

We obtained the data [12] from http://research.libd.org/spatialLIBD. We used the subject with brain ID Br5292, which contain 4 replicates, and in total includes 18,033 spots and 1,942 genes. The expression matrix was *log*-normalized and scaled to unit variance and zero mean.

### 2.6 Validation and Benchmarking

#### 2.6.1 Evaluation Metrics

Simulated Data In simulated datasets where the ground truth temporal progression of cells is known, we assessed the VAPOR’s inferred pseudotimes by computing the Pearson correlation coefficient between the ground truth pseudotimes and the process-specific pseudotime for each Ψ_*m*_. <u>Real Data</u> For real-world datasets, we utilized discrete known ordered labels, such as lineage cell types or time points. We converted these discrete labels into ordered variables. We compute Spearman’s rank correlation coefficient and Kendall’s tau coefficient.

#### 2.6.2 Benchmarking

We benchmarked VAPOR against Palantir[27] and Diffusion Pseudotime (DPT)[28]. For datasets where spliced and unspliced transcript counts are available, we also compare VAPOR to scVelo[19]. We use default parameters as recommended. When these methods require specifying an initial state or starting cell, we provide the ground truth starting cell in simulated dataset or use prior biological knowledge to select appropriate starting cells in real dataset.

## 3 Results

### 3.1 Simulation Study

To assess VAPOR’s performance in (1) capturing different gene expression dynamic patterns, and (2) decoupling co-occuring processes, we simulated three artificial gene expression datasets with a variety of gene dynamics, similar to the approach used in GeneTrajectory[24]. We simulated (1) a linear progression (**Fig. 2a**) in which genes sequentially reach their peak expression over time, (2) a cycling process (**Fig. 2b**) in which genes show periodic expression patterns over time; and (3) a mixture of two co-occurring processes (**Fig. 2c**), one process linear, and the other cyclic. In the last scenario, each cell state is determined by two independent variables - a pseudotime along the linear process and a pseudotime in the cyclic process.

**Figure 2:**
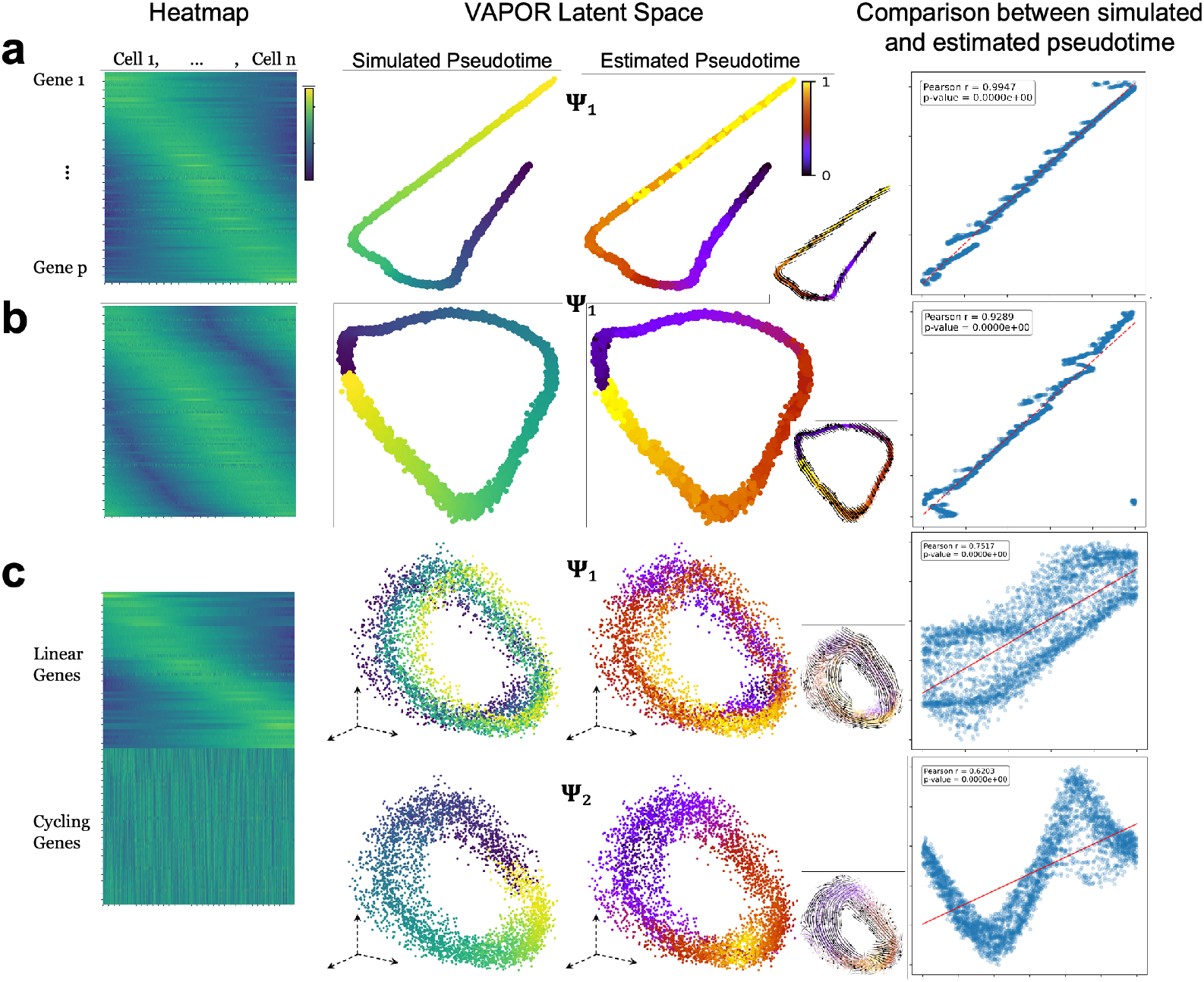
Simulation study. **a, Simulation of a linear progression process**. 4,000 cells and 100 genes; **b, Simulation of a cyclic process**. 4,000 cells and 100 genes; **c, Simulation of a combined linear and cyclic process**. 4,000 cells, 50 linear genes, 50 cycling genes. **First column:** Heatmaps of simulated gene expression data (rows: genes; columns: cells). **Second Column:** VAPOR’s latent representation of cells, colored by ground truth pseudotimes (linear process and cyclic process are visualized in 2-dimensions, the combined process is visualized in 3-dimensions). **Third column:** VAPOR’s latent representation of cells, colored by estimated pseudotimes from process-specific operators Ψ_*m*_. Bottom-right inset: Visualization of the latent subspace dynamics captured by each Ψ_*m*_, illustrating how VAPOR models each process separately. **Forth column:** Scatter plots comparing simulated (*x*-axis) with estimated (*y*-axis) pseudotimes.

As a result, VAPOR recapitulated the dynamic patterns of both the linear and cyclic processes, supported by Pearson correlations between ground truth and estimated pseudotimes (linear: *r* = 0.9947; cyclic *r* = 0.9289). In the scenario with coupled linear and cyclic processes, VAPOR decoupled the original mixture of two processes into two distinct representations. This is supported by the Pearson correlation coefficients *r* = 0.7517 with linear pseudotime inferred from Ψ_1_ and *r* = 0.6203 with cyclic pseudotime inferred from Ψ_2_.

We compared VAPOR’s performance with Palantir[27] and Diffusion Pseudotime (DPT)[28] (**Fig. S1**). For the linear process (**Fig. S1a, d**), VAPOR and Palantir performed comparably (Palantir: *r* = 0.9989; VAPOR: *r* = 0.9947), both nearly perfectly estimating the linear process pseudotime. In contrast, DPT’s estimation showed lower correlation (*r* = 0.1049) with the simulated pseudotime. For the cyclic process, VAPOR outperformed both Palantir[27] and DPT[28] (**Fig. S1b, d**). VAPOR’s estimation reached a high correlation of *r* = 0.9289 with the simulated pseudotime, while both Palantir and DPT’s estimations showed lower correlation (Palantir: *r* = 0.0155; DPT: *r* = −0.0330).

For the coupled process, Palantir and DPT each output a single pseudotime, which we used to compare against both the simulated linear pseudotime and cyclic pseudotime. VAPOR, decoupled the processes into two distinct ones (Ψ_1_ and Ψ_2_), allowed separate comparisons with the corresponding simulated process. For the linear process, Palantir (*r* = 0.8045) and DPT (*r* = 0.8903) performed better than VAPOR’s Ψ_1_ (*r* = 0.7517). For the cyclic process, however, VAPOR’s Ψ_2_ achieved higher correlation (*r* = 0.6203) compared to Palantir (*r* = 0.0990) and DPT (*r* = 0.1936) (**Fig. S1c, d**). These results demonstrate VAPOR’s advantage in decoupling and recapitulating distinct, co-occurring processes.

### 3.2 Decoupling Cell Differentiation and Maturation of Developmental Human Brain using scRNA-seq

We applied VAPOR to time-resolved scRNA-seq from the developing human cerebral cortex. VAPOR decoupled gene expression dynamics into two distinct processes: differentiation (Ψ_1_) and maturation (Ψ_2_), visualized on a UMAP embedding of VAPOR’s latent space (**Fig. 3a**). The differentiation process captured cell type transitions, showing clear progression from radial glia to intermediate progenitor cells and neurons; while the maturation process (Ψ_2_) captured age-related progression within cell types, suggesting how cells may develop specialized functions over time without changing their identity.

**Figure 3:**
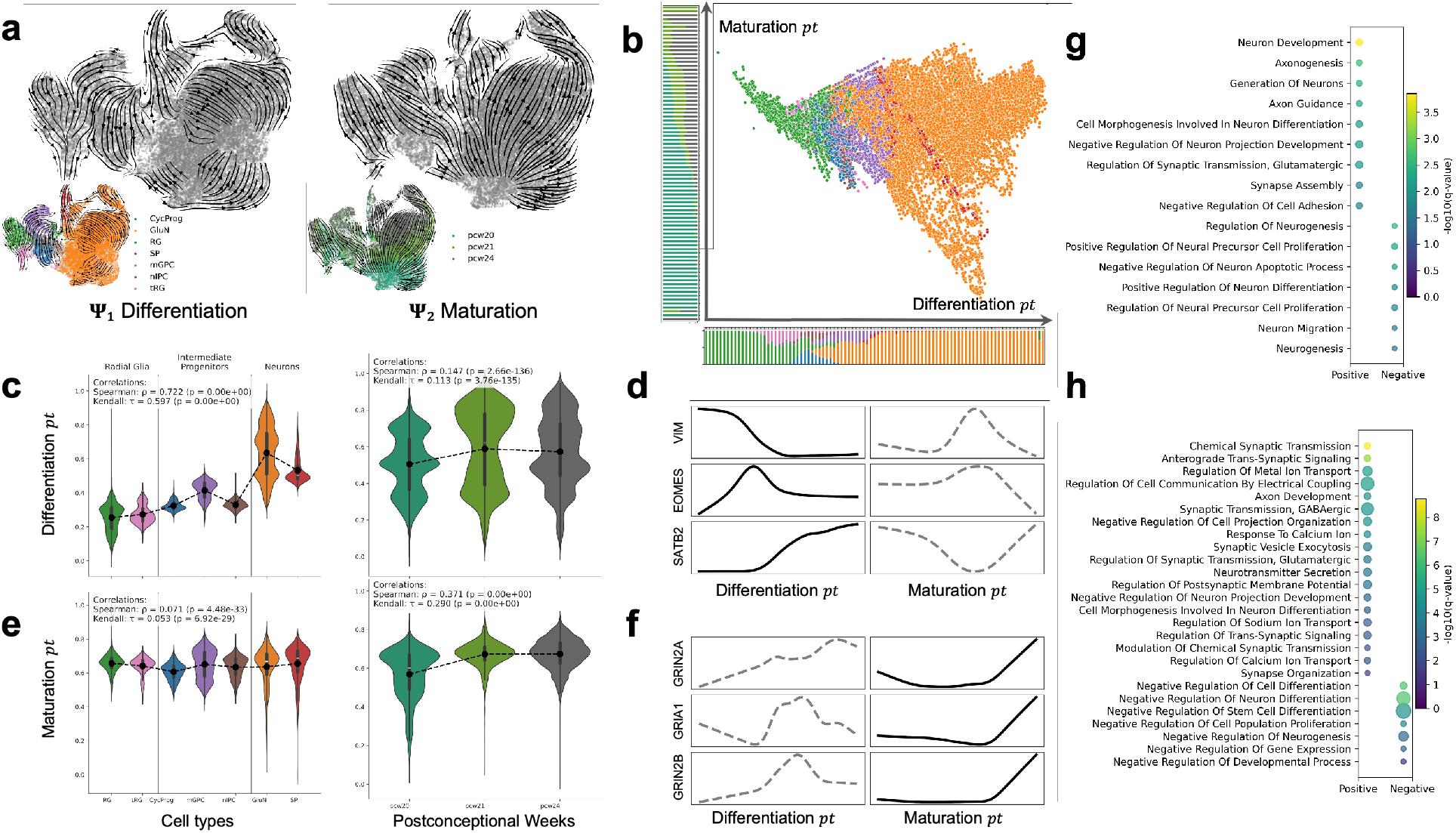
Decoupling Cell Differentiation and Maturation of Developmental Human Brain. **a, Streamplots of decoupled processes**. UMAP of developmental human brain scRNA-seq data (28,334 cells, 2,000 highly variable genes). Left: Ψ_1_ reveals cell type differentiation; Right: Ψ_2_ reveals age-dependent maturation. Insets are colored by cell type (RG: radial glia; tRG: truncated radial glia; cycProg: cycling progenitors; mGPC: multipotent glial progenitor cell; nIPC: neuronal intermediate progenitor cell; GluN: glutamatergic neuron; SP: subplate) and postconceptional weeks (pcw 20, 21, and 24). **b, Multifaceted pseudotimes Derived from** Ψ_1_ **and** Ψ_2_. Cells plotted according to differentiation pseudotime (from Ψ_1_, *x*-axis) and maturation pseudotime (from Ψ_2_, *y*-axis). Cell type composition is shown below the *x*-axis; age composition is shown to the left of the *y*-axis. **c, e, Violin plots of pseudotimes grouped by cell type and age. c**, Differentiation pseudotime (from Ψ_1_) and **e**, Maturation pseudotime (Ψ_2_): Left panels grouped by cell type; right panels grouped by age. **d, f, Gene expression trends along each decoupled process. d**, Expression of cell type marker genes (VIM, EOMES, SATB2) against differentiation pseudotime (left) and maturation pseudotime (right); **f**, Expression of maturation marker genes (GRIN2A, GRIN2B, GRIA1) against differentiation pseudotime (left) and maturation pseudotime (right). **g, h, Gene ontology (GO) enrichment analysis**. GO terms enriched among genes correlated with differentiation pseudotime (**g**) and maturation pseudotime(**h**).

To further characterize these processes, we derived pseudotimes for both Ψ_1_ and Ψ_2_ (**Fig. 3b**). Joint visualization of differentiation and maturation provides a complementary view of the multifaceted aspects of human brain development. Violin plots of pseudotimes across cell types and ages (**Fig. 3c,e**) further supported the distinction between these processes. Specifically, Ψ_1_ pseudotime (**Fig. 3c**) correlated strongly with differentiation lineage cell types (Spearman’s 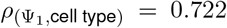, Spearman’s 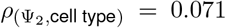), while Ψ_2_ pseudotime (**Fig. 3e**) correlated better with age (Spearman’s 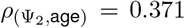, Spearman’s 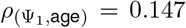). Compared to other methods (Palantir[27], DPT[28], and scVelo[19]) (**Fig. S2**), VAPOR showed comparable performance in capturing differentiation (VAPOR: Spearman’s *ρ* = 0.722; Palantir: Spearman’s *ρ* = 0.756; DPT: Spearman’s *ρ* = 0.754; scVelo: Spearman’s *ρ* = 0.743), and outperformed them in detecting age-associated dynamics(VAPOR: Spearman’s *ρ* = 0.371; Palantir: Spearman’s *ρ* = 0.057; DPT: Spearman’s *ρ* = 0.129; scVelo: Spearman’s *ρ* = 0.029).

We examined known marker genes along these pseudotimes (**Fig. 3d,f**) to demonstrate the distinction between differentiation and maturation. For instance, genes such as VIM (radial glia marker[29, 30]), EOMES (intermediate progenitors marker[30]), and SATB2 (glutamatergic neurons marker[31]) showed clear expression peaks at early, middle, and late stages, respectively, along the differntiation pseudotime. These trends were not observed along the maturation pseudotime (**Fig. 3d**). On the other hand, genes like GRIN2A, GRIN2B[32], and GRIA1[33] showed increasing expression along the maturation pseudotime, reflecting their roles in synaptic function and neuronal maturation (**Fig. 3f**). These patterns confirm that VAPOR effectively decouples these two critical aspects of brain development. Additionally, Gene Ontology (GO) enrichment analyses (with threshold FDR *q*-values *<* 0.05) provided differentiation-associated processes such as “Neuron Development”, “Cell Morphogenesis Involved in Neuron Differentiation” and “Regulation of Neurogenesis”(**Fig. 3g**) and maturation-associated processes like “Chemical Synaptic Transmission”, “Regulation of Metal Ion Transport”, “Neurotransmitter Secretion”, etc (**Fig. 3h**).

### 3.3 Capturing Inside-Out Formation Pattern of Human Cortical Layers using Spatial Transcriptomics

We applied VAPOR method to spatial transcriptomics data from the human dorsolateral prefrontal cortex, analyzing 1,942 genes across 18,033 spots from four replicates of subject Br5292. Despite not using spatial information during model training, our approach uncovered meaningful spatial patterns in cortical organization. Notably, the second decoupled process (Ψ_2_) by VAPOR aligned closely with the well-established ‘inside-out’ laminar structure of cortical layers (**Fig. 4a**). Visualization of Ψ_2_ on spatial coordinates showed streamlines progressing from deeper to upper cortical layers, confirming the layer-specific spatial organization (**Fig. 4b**). Violin plots of Ψ_2_ pseudotime across cortical layers (**Fig. 4c**) showed increasing values from deep to upper layers. Ψ_2_ pseudotime correlated with the laminar structure (Spearman’s *ρ* = 0.563).

**Figure 4:**
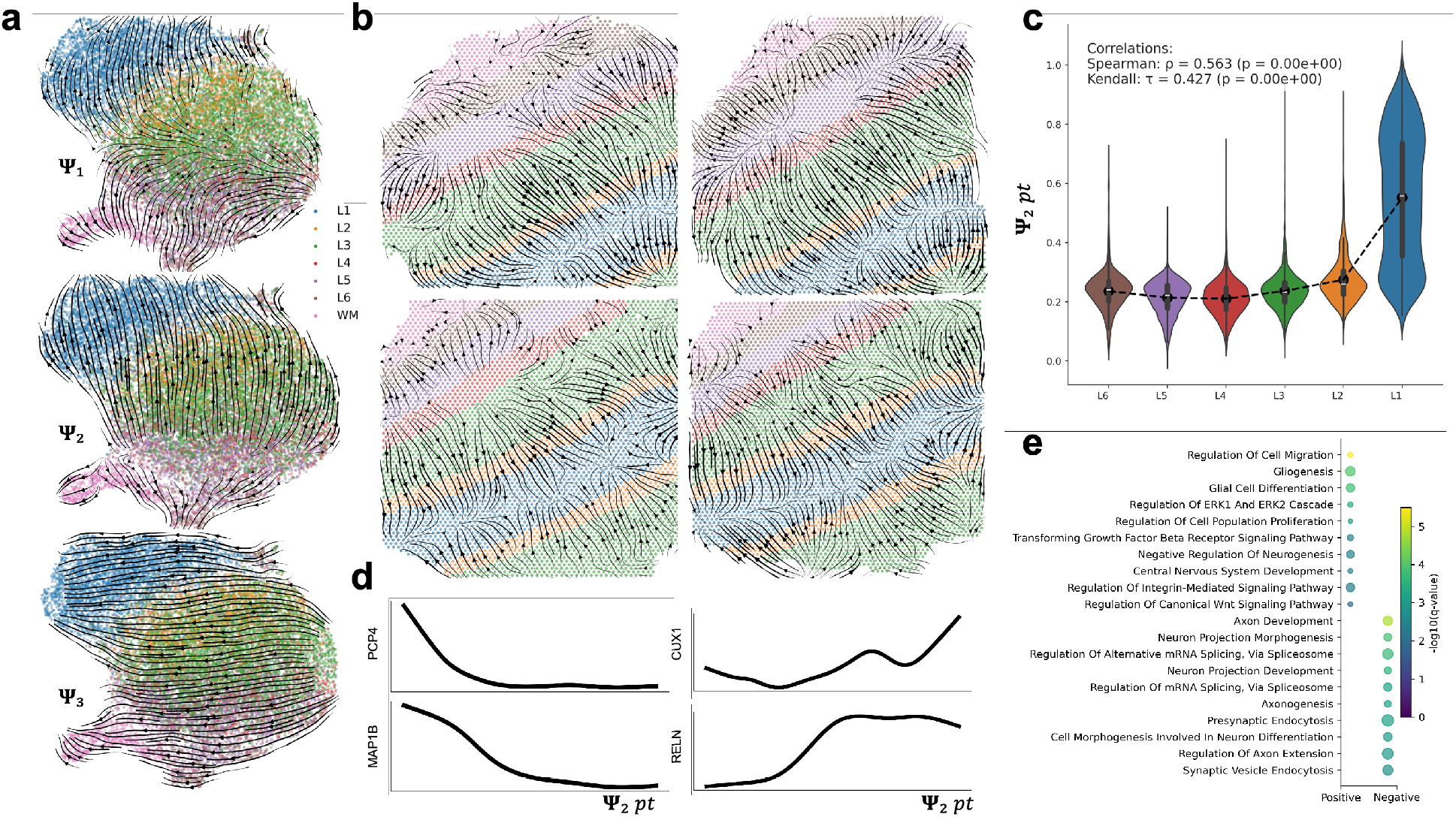
Capturing Inside-Out Formation Pattern of Human Cortical Layers using Spatial Transcriptomics. **a, Streamplots of decoupled processes on UMAP**. UMAP of spatial transcriptomics data from the human dorsolateral prefrontal cortex (Sample ID: Br5292, 4 replicates, 18,033 spots, 1,942 genes). Decoupled gene expression dynamics visualized on UMAP of the VAPOR’s latent space. From top to bottom: Ψ_1_,Ψ_2_, Ψ_3_. Spots are colored by cortical layers (L1-6) and white matter (WM). **b, Streamplots of** Ψ_2_ **on the spatial coordinates**. Streamplots of the dynamics captured by Ψ_2_ overlaid on the actual spatial coordinates of the tissue sections. Each panel represents a replicate of the sample. Spots are colored by cortical layers and white matter. **c, Violin plots of** Ψ_2_ **pseudotimes grouped by cortical layers. d, Expression of cortical specific marker genes**. PCP4, MAP1B, CUX1, and RELN against Ψ_2_ pseudotime. **e, Gene ontology (GO) enrichment analysis**. GO terms enriched among genes correlated with Ψ_2_ pseudotime(**g**).

This pattern was further supported by the expression trends of layer-specific marker genes along the Ψ_2_ pseudotime, including PCP4 (Layer V marker[34]), MAP1B (Neuronal migration marker[35]), CUX1 (Layers II-IV marker[36]), RELN (Layer I marker[37]) (**Fig. 4d**). The gradual changes in the expression of these genes reflect the transition from deeper to upper layers.

Gene Ontology enrichment analysis (with threshold FDR *q*-values *<* 0.05) of the genes correlated with Ψ_2_ pseudotime highlighted processes crucial to layer-specific functions, such as “Regulation of Cell Migration”, “ERK1/ERK2 Cascade” (**Fig. 4e**). Additional processes identified by VAPOR revealed complementary aspects of cortical organization. Specifically, Ψ_1_ **(Fig. S3a)** was enriched for myelination and axon development terms, suggesting white matter-related processes, and Ψ_3_ (**Fig. S3b**) showed enrichment in neuronal projection and synaptic organization terms, potentially indicating patterns related to neural circuit. These findings demonstrate VAPOR’s ability to decouple biologically meaningful dynamics, especially capturing layer-specific organization without relying on prior spatial information.

## 4 Discussion

VAPOR is a novel computational method to decouple co-occurring biological processes in development and reveal underlying gene expression dynamics associate with each process from single-cell and spatial transcriptomics data. Particularly, VAPOR combines a variational autoencoder model with transport operators to model gene expression dynamics. The key to VAPOR’s efficacy in capturing multiple, co-occurring dynamics is the decomposition of VAE’s latent subspaces, each modeled by its own ordinary differential equation. VAPOR learns multifaceted development in a fully data-driven, unsupervised manner with only limited assumptions about the underlying biology. In contrast to previous methods that simplify development to a single pseudotime axis or assume uniform kinetic rates, VAPOR can capture and decouple multiple, co-occurring processes, potentially revealing their contributions to the overall development.

In this work, we have demonstrated VAPOR’s capacities to decoupling both simulated and real-world data, including developmental human brain scRNA-seq data and human dorsolateral prefrontal cortex spatial transcriptomics data. However, our method can be generalized to other single-cell or spatial modalities, including but not limited to scATAC-seq and spatial epigenomics. We anticipate VAPOR can be applied to spatial time-resolved data to elucidate spatio-temporal dynamics in development[13]. We also anticipate VAPOR’s framework can be extended to integrate multi-omics data, capturing the interplay between epigenetic and transcriptomic dynamics underlying co-occurring biological processes and elucidating multi-layered molecular mechanisms driving them.

Finally, VAPOR currently has several limitations. First, the model relies solely on feature (gene) similarities, without incorporating temporal or spatial context into the training process. This may limit VAPOR’s accuracy in modeling complex development. Second, the inferred direction of progression is bidirectional, which leads to ambiguous interpretation of the developmental progression. We may address these limitations in the future by incorporating prior knowledge such as temporal and spatial proximity or known starting points into the model.

## 5 Code availability

Python implementations of VAPOR are available on GitHub (https://github.com/daifengwanglab/VAPOR).

## 6 Acknowledgment

This work is supported in part by National Institutes of Health grants, RF1MH128695, R01AG067025, R21NS128761, R21NS127432, P50HD105353, and National Science Foundation Career Award 2144475.

## S1 Supplementary Figures

**Figure S1:**
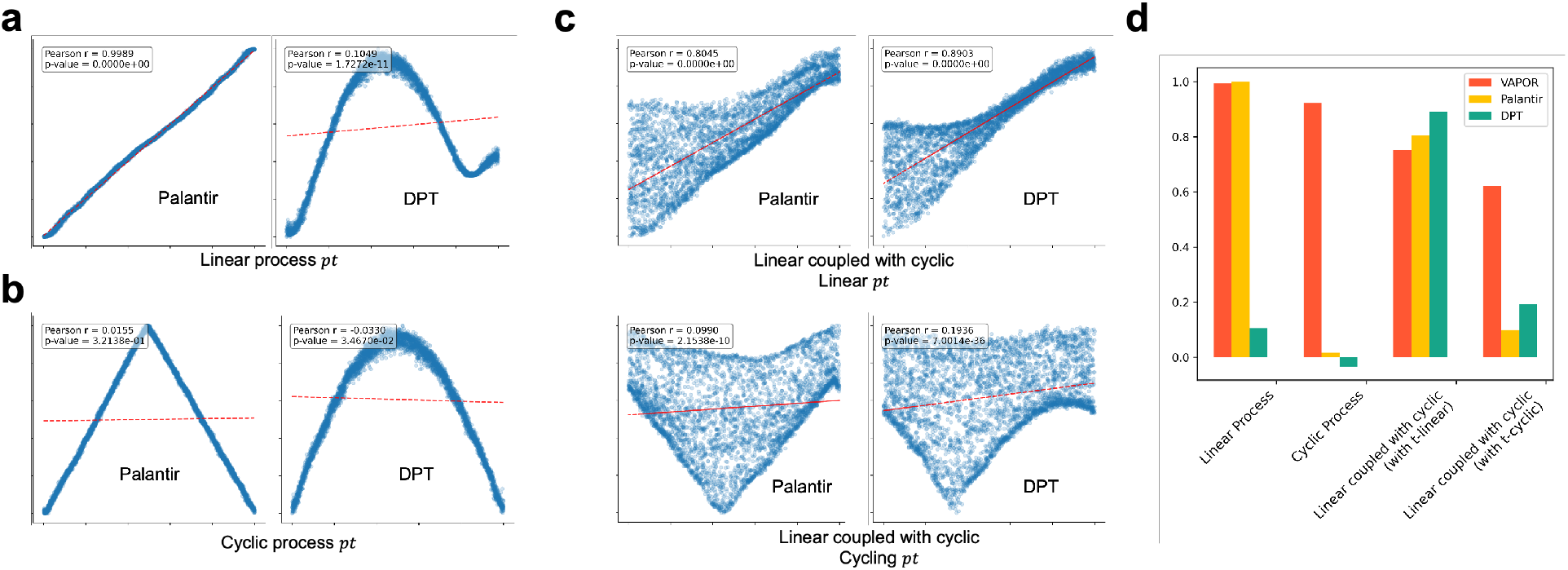
Benchmarking on simulation data. **a-c**, Scatter plots comparing simulated (*x*-axis) versus estimated (*y*-axis) pseudotimes for Palantir (left) and DPT (right). Results are shown for **a**, linear process; **b**, cyclic process; **c**. coupled process. **d**. Pearson corelation coefficients for VAPOR, Palantir, and DPT across different simulated data.

**Figure S2:**
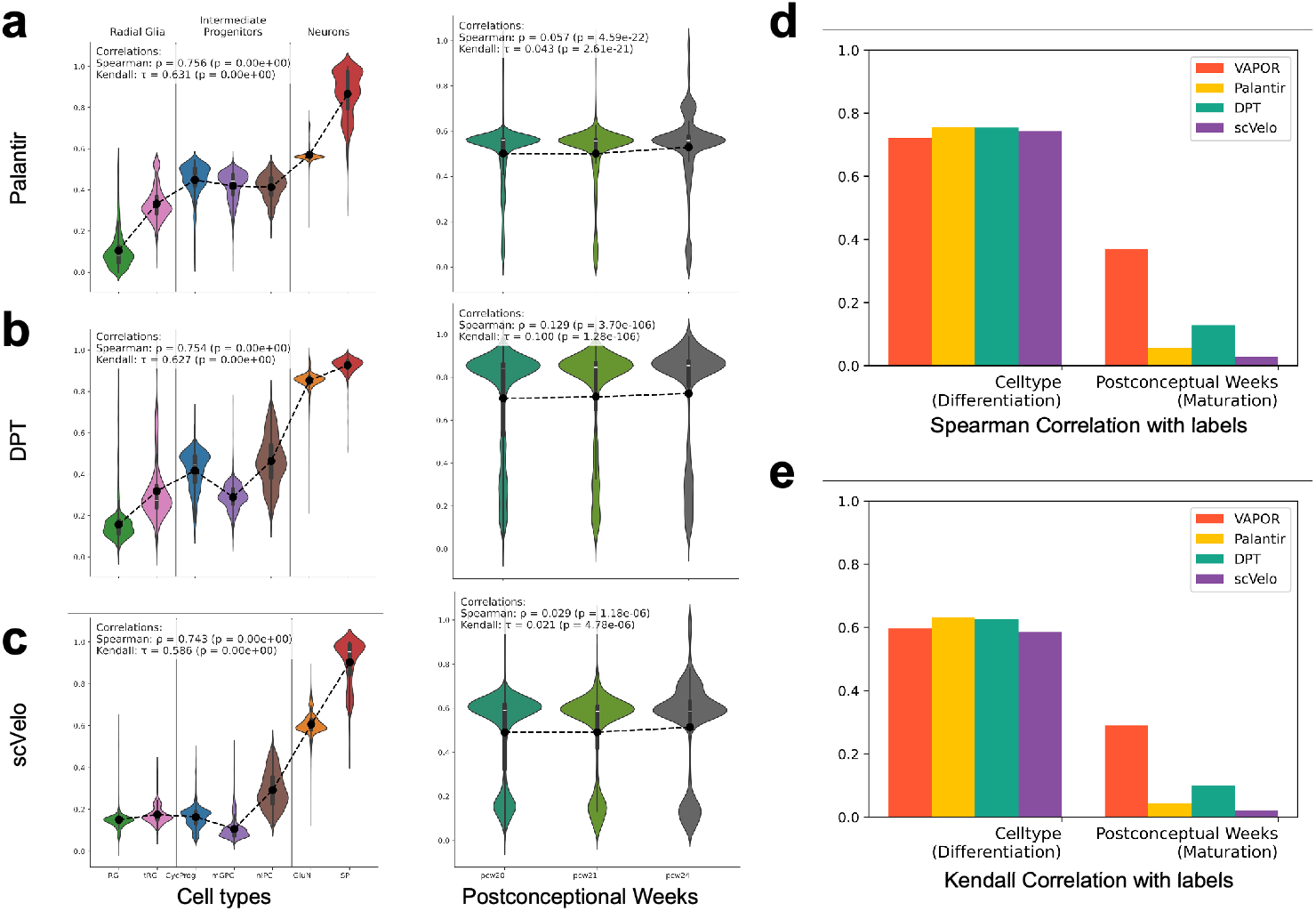
Benchmarking on developmental human brain scRNA-seq data. **a-c**, Violin plots of pseudotimes grouped by cell type and age. **a**, Palantir; **b**, DPT; **c**, scVelo. Left panels grouped by cell type; right panels grouped by age. **d**, Spearman corelation coefficients for VAPOR, Palantir, DPT, and scVelo when comparing inferred pseudotimes with known sequential ordered cell types, and ages. **e**, Kendall rank corelation coefficients for VAPOR, Palantir, DPT, and scVelo when comparing inferred pseudotimes with known sequential ordered cell types, and ages.

**Figure S3:**
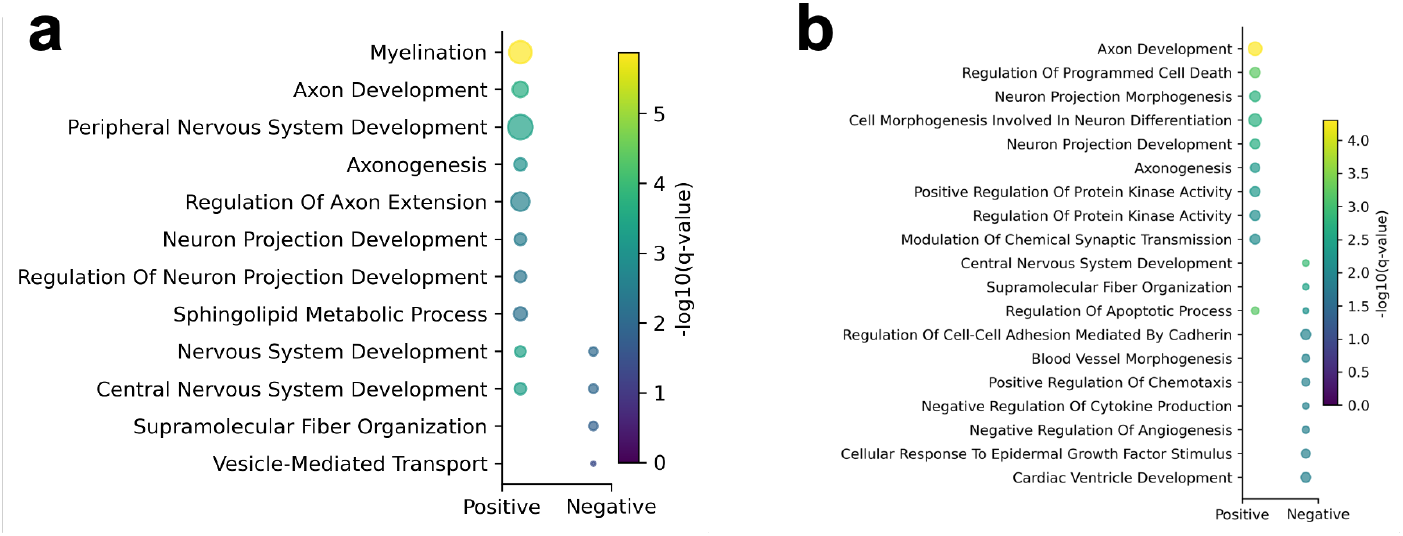
Gene ontology (GO) enrichment analysis for spatial transcriptomics data. **a**, GO terms enriched among genes correlated with Ψ_1_ pseudotime. **b**, GO terms enriched among genes correlated with Ψ_3_ pseudotime.

## References

[1] Eric H Davidson, Jonathan P Rast, Paola Oliveri, Andrew Ransick, Cristina Calestani, Chiou-Hwa Yuh, Takuya Minokawa, Gabriele Amore, Veronica Hinman, Cesar Arenas-Mena, et al. A genomic regulatory network for development. science, 295(5560):1669–1678, 2002.

[2] Michael Levine and Eric H Davidson. Gene regulatory networks for development. Proceedings of the National Academy of Sciences, 102(14):4936–4942, 2005.

[3] Albert-Laszlo Barabasi and Zoltan N Oltvai. Network biology: understanding the cell’s functional organization. Nature reviews genetics, 5(2):101–113, 2004.

[4] Amit Tzur, Ran Kafri, Valerie S LeBleu, Galit Lahav, and Marc W Kirschner. Cell growth and size homeostasis in proliferating animal cells. Science, 325(5937):167–171, 2009.

[5] Cédric Blanpain and Benjamin D Simons. Unravelling stem cell dynamics by lineage tracing. Nature reviews Molecular cell biology, 14(8):489–502, 2013.

[6] Lisette Hari, Iris Miescher, Olga Shakhova, Ueli Suter, Lynda Chin, Makoto Taketo, William D Richardson, Nicoletta Kessaris, and Lukas Sommer. Temporal control of neural crest lineage generation by wnt/β-catenin signaling. Development, 139(12):2107–2117, 2012.

[7] Disha Sood, Dana M Cairns, Jayanth M Dabbi, Charu Ramakrishnan, Karl Deisseroth, Lauren D Black III, Sabato Santaniello, and David L Kaplan. Functional maturation of human neural stem cells in a 3d bioengineered brain model enriched with fetal brain-derived matrix. Scientific reports, 9(1):17874, 2019.

[8] Alexandro E Trevino, Fabian Mü ller, Jimena Andersen, Laksshman Sundaram, Arwa Kathiria, Anna Shcherbina, Kyle Farh, Howard Y Chang, Anca M Paşca, Anshul Kundaje, et al. Chromatin and gene-regulatory dynamics of the developing human cerebral cortex at single-cell resolution. Cell, 184(19):5053–5069, 2021.

[9] Markus Mittnenzweig, Yoav Mayshar, Saifeng Cheng, Raz Ben-Yair, Ron Hadas, Yoach Rais, Elad Chomsky, Netta Reines, Anna Uzonyi, Lior Lumerman, et al. A single-embryo, single-cell time-resolved model for mouse gastrulation. Cell, 184(11):2825–2842, 2021.

[10] Emelie Braun, Miri Danan-Gotthold, Lars E Borm, Ka Wai Lee, Elin Vinsland, Peter Lönnerberg, Lijuan Hu, Xiaofei Li, Xiaoling He, Žaneta Andrusivová, et al. Comprehensive cell atlas of the first-trimester developing human brain. Science, 382(6667):eadf1226, 2023.

[11] Nicola Micali, Shaojie Ma, Mingfeng Li, Suel-Kee Kim, Xoel Mato-Blanco, Suvimal Kumar Sindhu, Jon I Arellano, Tianliuyun Gao, Mikihito Shibata, Kevin T Gobeske, et al. Molecular programs of regional specification and neural stem cell fate progression in macaque telencephalon. Science, 382(6667):eadf3786, 2023.

[12] Kristen R Maynard, Leonardo Collado-Torres, Lukas M Weber, Cedric Uytingco, Brianna K Barry, Stephen R Williams, Joseph L Catallini, Matthew N Tran, Zachary Besich, Madhavi Tippani, et al. Transcriptome-scale spatial gene expression in the human dorsolateral prefrontal cortex. Nature neuroscience, 24(3):425–436, 2021.

[13] Di Zhang, Leslie A Rubio Rodriguez-Kirby, Yingxin Lin, Mengyi Song, Li Wang, Lijun Wang, Shigeaki Kanatani, Tony Jimenez-Beristain, Yonglong Dang, Mei Zhong, et al. Spatial dynamics of mammalian brain development and neuroinflammation by multimodal tri-omics mapping. bioRxiv, pages 2024–07, 2024.

[14] Meng Zhang, Xingjie Pan, Won Jung, Aaron R Halpern, Stephen W Eichhorn, Zhiyun Lei, Limor Cohen, Kimberly A Smith, Bosiljka Tasic, Zizhen Yao, et al. Molecularly defined and spatially resolved cell atlas of the whole mouse brain. Nature, 624(7991):343–354, 2023.

[15] Cole Trapnell, Davide Cacchiarelli, Jonna Grimsby, Prapti Pokharel, Shuqiang Li, Michael Morse, Niall J Lennon, Kenneth J Livak, Tarjei S Mikkelsen, and John L Rinn. The dynamics and regulators of cell fate decisions are revealed by pseudotemporal ordering of single cells. Nature biotechnology, 32(4):381–386, 2014.

[16] F Alexander Wolf, Fiona K Hamey, Mireya Plass, Jordi Solana, Joakim S Dahlin, Berthold Göttgens, Nikolaus Rajewsky, Lukas Simon, and Fabian J Theis. Paga: graph abstraction reconciles clustering with trajectory inference through a topology preserving map of single cells. Genome biology, 20:1–9, 2019.

[17] Kelly Street, Davide Risso, Russell B Fletcher, Diya Das, John Ngai, Nir Yosef, Elizabeth Purdom, and Sandrine Dudoit. Slingshot: cell lineage and pseudotime inference for single-cell transcriptomics. BMC genomics, 19:1–16, 2018.

[18] Gioele La Manno, Ruslan Soldatov, Amit Zeisel, Emelie Braun, Hannah Hochgerner, Viktor Petukhov, Katja Lidschreiber, Maria E Kastriti, Peter Lönnerberg, Alessandro Furlan, et al. Rna velocity of single cells. Nature, 560(7719):494–498, 2018.

[19] Volker Bergen, Marius Lange, Stefan Peidli, F Alexander Wolf, and Fabian J Theis. Generalizing rna velocity to transient cell states through dynamical modeling. Nature biotechnology, 38(12):1408–1414, 2020.

[20] Mingze Gao, Chen Qiao, and Yuanhua Huang. Unitvelo: temporally unified rna velocity reinforces single-cell trajectory inference. Nature Communications, 13(1):6586, 2022.

[21] Adam Gayoso, Philipp Weiler, Mohammad Lotfollahi, Dominik Klein, Justin Hong, Aaron Streets, Fabian J Theis, and Nir Yosef. Deep generative modeling of transcriptional dynamics for rna velocity analysis in single cells. Nature methods, 21(1):50–59, 2024.

[22] Volker Bergen, Ruslan A Soldatov, Peter V Kharchenko, and Fabian J Theis. Rna velocity—current challenges and future perspectives. Molecular systems biology, 17(8):e10282, 2021.

[23] Cheng Frank Gao, Suriyanarayanan Vaikuntanathan, and Samantha J Riesenfeld. Dissection and integration of bursty transcriptional dynamics for complex systems. Proceedings of the National Academy of Sciences, 121(18):e2306901121, 2024.

[24] Rihao Qu, Xiuyuan Cheng, Esen Sefik, Jay S Stanley III, Boris Landa, Francesco Strino, Sarah Platt, James Garritano, Ian D Odell, Ronald Coifman, et al. Gene trajectory inference for single-cell data by optimal transport metrics. Nature Biotechnology, pages 1–11, 2024.

[25] Diederik P Kingma. Auto-encoding variational bayes. arXiv preprint arXiv:1312.6114, 2013.

[26] Benjamin Culpepper and Bruno Olshausen. Learning transport operators for image manifolds. Advances in neural information processing systems, 22, 2009.

[27] Manu Setty, Vaidotas Kiseliovas, Jacob Levine, Adam Gayoso, Linas Mazutis, and Dana Pe’Er. Characterization of cell fate probabilities in single-cell data with palantir. Nature biotechnology, 37(4):451–460, 2019.

[28] Laleh Haghverdi, Maren Büttner, F Alexander Wolf, Florian Buettner, and Fabian J Theis. Diffusion pseudotime robustly reconstructs lineage branching. Nature methods, 13(10):845–848, 2016.

[29] Alex A Pollen, Tomasz J Nowakowski, Jiadong Chen, Hanna Retallack, Carmen Sandoval-Espinosa, Cory R Nicholas, Joe Shuga, Siyuan John Liu, Michael C Oldham, Aaron Diaz, et al. Molecular identity of human outer radial glia during cortical development. Cell, 163(1):55–67, 2015.

[30] Chris Englund, Andy Fink, Charmaine Lau, Diane Pham, Ray AM Daza, Alessandro Bulfone, Tom Kowalczyk, and Robert F Hevner. Pax6, tbr2, and tbr1 are expressed sequentially by radial glia, intermediate progenitor cells, and postmitotic neurons in developing neocortex. Journal of Neuroscience, 25(1):247–251, 2005.

[31] Dino P Leone, Whitney E Heavner, Emily A Ferenczi, Gergana Dobreva, John R Huguenard, Rudolf Grosschedl, and Susan K McConnell. Satb2 regulates the differentiation of both callosal and subcerebral projection neurons in the developing cerebral cortex. Cerebral cortex, 25(10):3406–3419, 2015.

[32] Pierre Paoletti, Camilla Bellone, and Qiang Zhou. Nmda receptor subunit diversity: impact on receptor properties, synaptic plasticity and disease. Nature Reviews Neuroscience, 14(6):383–400, 2013.

[33] Jeremy M Henley and Kevin A Wilkinson. Ampa receptor trafficking and the mechanisms underlying synaptic plasticity and cognitive aging. Dialogues in clinical neuroscience, 15(1):11–27, 2013.

[34] Hongkui Zeng, Elaine H Shen, John G Hohmann, Seung Wook Oh, Amy Bernard, Joshua J Royall, Katie J Glattfelder, Susan M Sunkin, John A Morris, Angela L Guillozet-Bongaarts, et al. Large-scale cellular-resolution gene profiling in human neocortex reveals species-specific molecular signatures. Cell, 149(2):483–496, 2012.

[35] Junlin Teng, Yosuke Takei, Akihiro Harada, Takao Nakata, Jianguo Chen, and Nobutaka Hirokawa. Synergistic effects of map2 and map1b knockout in neuronal migration, dendritic outgrowth, and microtubule organization. The Journal of cell biology, 155(1):65–76, 2001.

[36] Marta Nieto, Edwin S Monuki, Hua Tang, Jaime Imitola, Nicole Haubst, Samia J Khoury, Jim Cunningham, Magdalena Gotz, and Christopher A Walsh. Expression of cux-1 and cux-2 in the subventricular zone and upper layers ii–iv of the cerebral cortex. Journal of Comparative Neurology, 479(2):168–180, 2004.

[37] Gundela Meyer, Carlos Gustavo Perez-Garcia, Hajnalka Abraham, and Daniel Caput. Expression of p73 and reelin in the developing human cortex. Journal of Neuroscience, 22(12):4973–4986, 2002.

